# Identification of a new human polyomavirus in distinct populations and tissues

**DOI:** 10.1101/2023.06.22.545219

**Authors:** Carolina Torres, Rita M. Correa, María A. Picconi, Christopher B. Buck, Viviana A. Mbayed

**Affiliations:** Universidad de Buenos Aires, Facultad de Farmacia y Bioquímica, Instituto de Investigaciones en Bacteriología y Virología Molecular (IBaViM), Buenos Aires, Argentina; Consejo Nacional de Investigaciones Científicas y Técnicas (CONICET), Buenos Aires, Argentina; Servicio Virus Oncogénicos, Departamento de Virología, Instituto Nacional de Enfermedades Infecciosas - ANLIS “Dr. Carlos G. Malbrán,” Buenos Aires, Argentina; Lab of Cellular Oncology, National Cancer Institute, NIH, Bethesda, MD, USA

## Abstract

**Objectives:** This work aimed to characterize a novel human polyomavirus (HPyV) with cutaneous tropism.

**Methods:** Swabs of healthy skin (forehead) of 75 immunocompetent individuals from Argentina were screened for HPyV through sequence amplification techniques. Publicly available metagenomic datasets were also analyzed.

**Results:** A previously unknown polyomavirus sequence was detected in two skin swab samples. A nearly identical sequence was detected in public datasets representing metagenomic surveys of human skin and feces. Further analyses showed that the new polyomavirus diverges from its nearest relative, human polyomavirus 6 (HPyV6), by 17.3-17.7% (in nucleotides for the large T antigen), which meets criteria for a new species designation in the genus *Deltapolyomavirus*.

The screening also revealed more distant HPyV6 relatives in macaque genital and chimpanzee fecal datasets. Since polyomaviruses are generally thought to cospeciate with mammalian hosts, the high degree of similarity to HPyV6 suggests the new polyomavirus species is human-tropic.

**Conclusions:** A novel polyomavirus was identified and characterized from samples of distinct populations and tissues. We suggest the common name human polyomavirus 16 (HPyV16).

## INTRODUCTION

Several human polyomaviruses (HPyVs) have been described in the last 15 years, including some with cutaneous tropism, such as the Merkel cell carcinoma-associated polyomavirus (MCPyV), trichodysplasia spinulosa-associated polyomavirus (TSPyV) and human polyomaviruses 6 (HPyV6) and 7 (HPyV7) [1–3]. In this report, we identified a previously unknown HPyV species in healthy skin samples from Argentina and in metagenomic datasets representing skin and fecal samples from the United States and Finland.

## METHODS

### Clinical samples

In the framework of a cross-sectional study carried out to analyze the presence of human polyomaviruses with cutaneous tropism in Argentina, healthy skin samples (forehead swabs) of immunocompetent individuals (n=75) collected in 2014-2015 were analyzed. The study was carried out following The Code of Ethics of the World Medical Association (Declaration of Helsinki) and was approved by the institutional Ethics Committee: “Comité de Ética en Investigaciones” de la Administración Nacional de Laboratorios e Institutos de Salud (ANLIS).

### Sample processing and sequencing

Swabs were immersed in 1 ml of saline solution, nucleic acids were extracted (QIAmp DNA Mini Kit, QIAGEN) and a multiplex nested PCR was implemented to detect known skin-tropic HPyVs (MCPyV, TSPyV, HPyV6 and HPyV7), based on a previous design [4]. Sequencing of PCR amplicons revealed the presence of two cases of a divergent HPyV, genetically close to HPyV6.

To obtain the full-length genome of the HPyV6-like sequence, a rolling circle amplification (RCA) with HPyV6-specific primers was performed in the two positive samples (sample 1 and sample 2), based on a similar strategy previously reported [5] but using the Phi29 DNA Polymerase (Thermo Scientific) and newly designed primers (Table S1). The RCA reaction was followed by two nested PCRs to cover the complete genome of the new virus (Table S1). Only one of the divergent HPyV genomes (isolate 39H, from sample 1) could be amplified and fully sequenced using the Sanger method (Macrogen, Korea).

### Detection of HPyV16 in public databases

Diamond 2.0 software was used to scan Sequence Read Archive (SRA) records for sequences resembling the helicase domain of polyomavirus Large T antigen, with settings --block-size 7 --index-chunks 1 --evalue 0.00001 --outfmt 6 qseqid sseqid evalue qseq [6]. Megahit 1.2.9 was used for *de novo* assembly of SRA records of interest, with default settings [7]. Contigs of interest were inspected and polished using CLC Genomics Workbench 22.

### Genomic characterization

Annotations of the early and late open reading frames (ORFs) were made in reference to the HPyV6 genome [2]. The identification of the large T (LT) splice donor and acceptor sites was carried out by searching for canonical splice donor and acceptor sites followed by a selection of those that are well conserved between the new HPyV and HPyV6, for which splice sites were experimentally determined [8]. Rating values were calculated with the Human Splice Finder tool [9].

### Genetic divergence and phylogenetic analyses

Nucleotide and amino acid pairwise identity between the new HPyV and members of the *Deltapolyomavirus* genus were estimated for the LT coding region using SDT v.1.2 [10].

Phylogenetic analyses were performed on the complete genome and LT amino acid sequences using Maximum likelihood and Bayesian methods. Maximum Likelihood trees were obtained using IQ-TREE v1.6, with an appropriate substitution model estimated with ModelFinder. Confidence was evaluated with the SH approximate likelihood ratio test (SH-aLRT) (1,000 replicates) and the Ultrafast Bootstrap method (UFB) (10,000 replicates). The majority-rule consensus trees were obtained from Bayesian analyses performed on MrBayes v3.2.7. Analyses were run until convergence, evaluated through effective sampling sizes higher than 200, and 10% was discarded as burn-in.

## RESULTS

A new human-associated polyomavirus was detected in two skin swab samples from Argentina (2/75, 2.7%) and in public databases from metagenomic studies of human skin (NCBI SRA accession number SRR3644061) and feces (SRR6914032) from the United States and Finland, respectively. The new sequence was not detected in Diamond screens of several hundred thousand SRA datasets representing a wide range of non-human animals. The screening did, however, reveal two more distant HPyV6 relatives in macaque genital (SRR8173832) and chimpanzee fecal (SRR8205461) samples.

The candidate sixteenth human-associated polyomavirus (HPyV16, Table S2) is 85% identical to its nearest known relative, HPyV6, at the whole genome nucleotide level. It displays the typical genomic organization of polyomaviruses: a circular double-stranded DNA genome, an early region encoding the LT and small T (sT) antigens, and a late region encoding the structural viral proteins on opposite strands, with a regulatory region separating both regions (Figure 1, Table S3).

**Figure 1.**
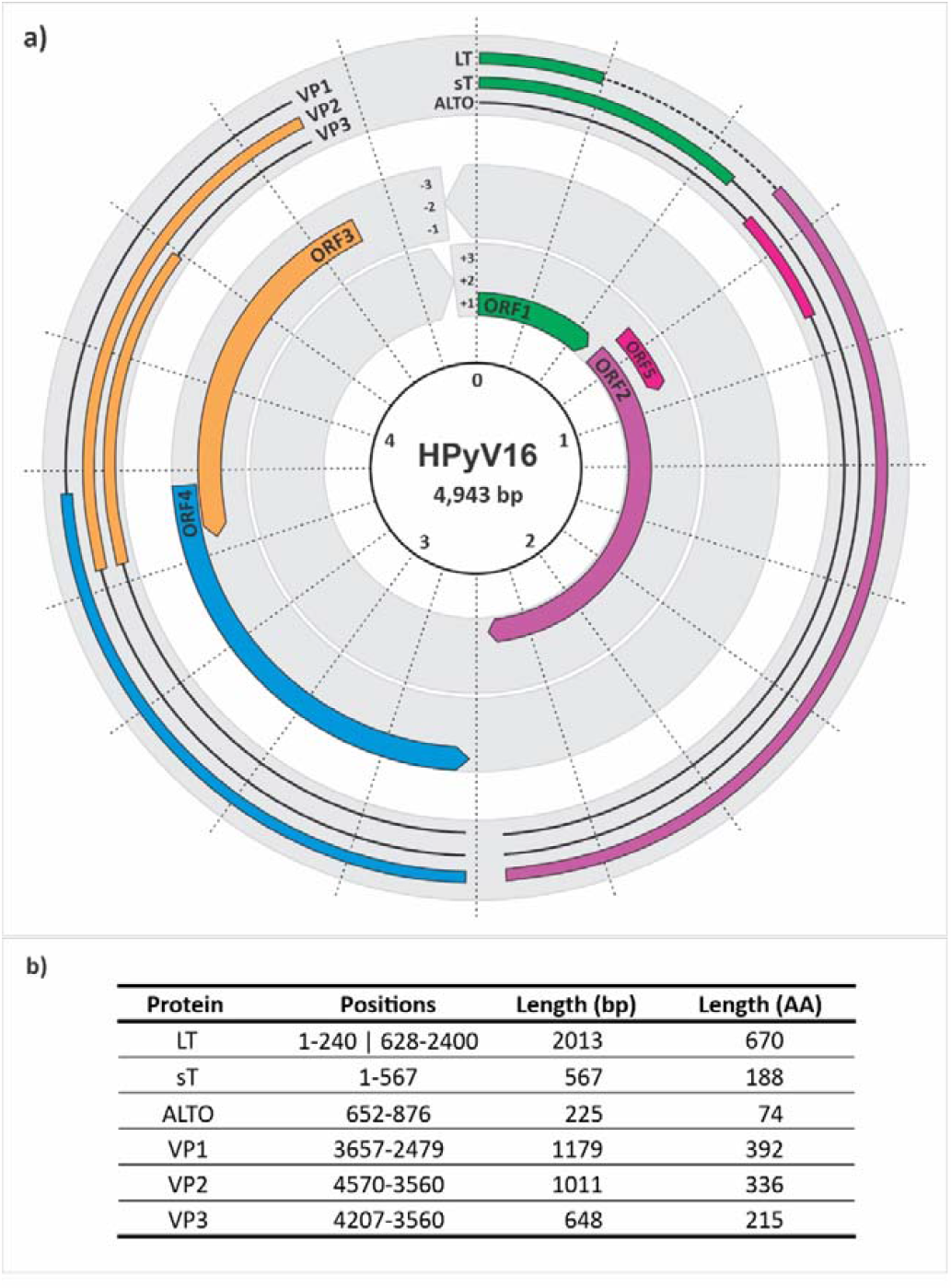
**(a)** Predicted genome organization, transcripts and proteins of the HPyV16. The ORFs are indicated in the inner layer and the transcripts are shown in the outer layer. The LT intron is shown as a dashed line. Radial dotted lines show 250 basepair (bp) increments. The regulatory region, located between early and late regions, encompasses 373 bp. The size of the genome corresponds to the two viruses recovered from skin, whereas that detected in feces has five nucleotides more in the intronic region of LT and sT. **(b)** Predicted proteins.

The three HPyV16 isolates characterized in this work clustered together in the phylogenetic trees (Figure 2 and Figure S1-S3) and showed pairwise identity in the coding LT region >96% in nucleotides and amino acids (Table S4 and Table S5), being more closely related those two complete genomes detected in the skin (Figure 2, Figure S1-S3). The new sequences diverge by 17.3-17.7% (nucleotide identity) from HPyV6 in the LT coding region (Table S4).

**Figure 2.**
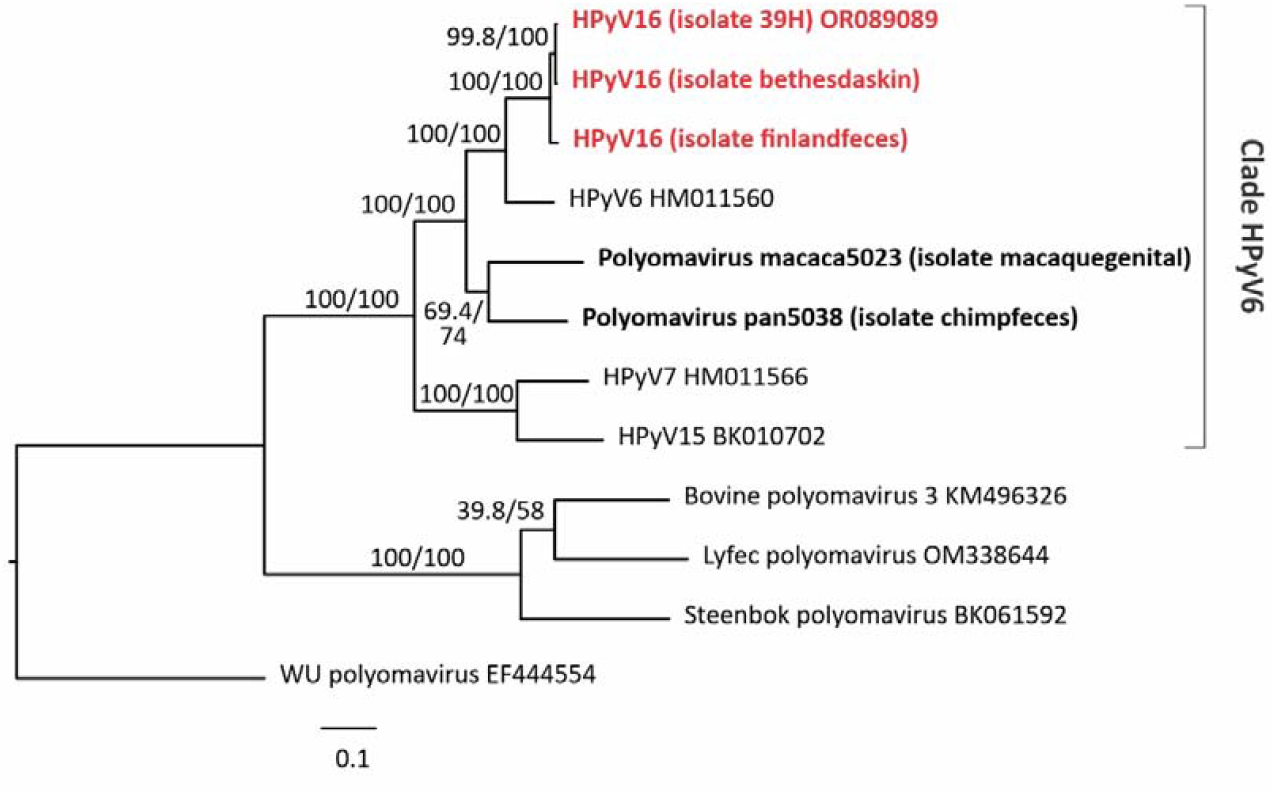
Maximum Likelihood phylogenetic tree based on complete genome sequences of the HPyV16 (in bold and red) and other members of the HPyV6 clade in the Wuki clade. Wu polyomavirus was used as outgroup. The two distant HPyV6 relatives detected in macaque genital (isolate: macaquegenital) and chimpanzee fecal (isolate: chimpfeces) samples are also included (in bold). The analysis was performed under the K3Pu+F+R3 nucleotide substitution model, estimated with ModelFinder in IQ-TREE. The SH-aLRT/UFB values are indicated at nodes.

Phylogenetic analyses show that HPyV16 occupies a discrete clade formed by HPyVs 6, 7, and 15. The clade also includes the two polyomaviruses detected in macaque and chimpanzee samples (Figure 2 and Figures S1-S3). The primate HPyV6 clade is distinct from a related polyomavirus clade associated with artiodactylans [11].

The second sample from Argentina, from which only a small sequence fragment could be obtained (isolate 57H), showed higher identity with the strain identified in the metagenome from feces (1 substitution in 165 nts) than with the other detected in the skin (3 or 4 substitutions) (Figure S4).

## DISCUSSION

During screening for human polyomaviruses in healthy skin swabs of immunocompetent adults from Argentina, a previously unknown polyomavirus sequence was found. In parallel to these findings, other two nearly identical viral genomes were assembled from metagenomic surveys available via public databases.

Further analysis of the viral genomes revealed that they grouped within the *Deltapolyomavirus* genus, being phylogenetically close to HPyV6, but fitting the criteria to be considered as a new viral species, for which the common name “human polyomavirus 16” (HPyV16) is proposed.

The classification criteria included: information on the natural host, observed genetic distance >15% for the LT coding nucleotide sequence relative to the most closely related known species, and unambiguous phylogenetic placement in analyses of LT amino acid sequences [12,13]. These criteria are fulfilled for the new sequences described in this study.

It is worth noting that the two HPyV16 sequences found in skin samples from Argentina and the USA are genetically closer to each other than to those found in feces from Finland. The Finland sequence was similar to that obtained in a small fragment from a third skin sample from Argentina. As seen in other HPyVs shed in skin and feces, such as MCPyV, sequences did not cluster based on the tissue or biological sample where the virus was found, but based on population and geography [14]. Further research is needed to reveal whether this pattern is also followed by HPyV16.

Finally, the report of unknown viruses allows a better understanding of human virome and is the basis for future studies focused on elucidating their potential role in human diseases and, potentially, vaccine development [15].

## DATA ACCESS

The complete genome sequences reported in this work have been deposited in GenBank under the accession numbers OR089089, XXXXXXXX – XXXXXXXX.

## AUTHOR CONTRIBUTIONS

CT designed and performed the experiments, analyzed data, acquired financial support, and drafted the article. RMC collected samples, performed the experiments, and provided critical revision of the article. MAP performed the experiments, analyzed data, acquired financial support, and provided critical revision of the article. CBB performed experiments, analyzed data, and edited article text and figures. VAM analyzed data, acquired financial support, and provided critical revision of the article. All author approved the final version of the manuscript.

