## Supplementary Figures S1-S4 for "Identification of a new human polyomavirus in distinct populations and tissues"

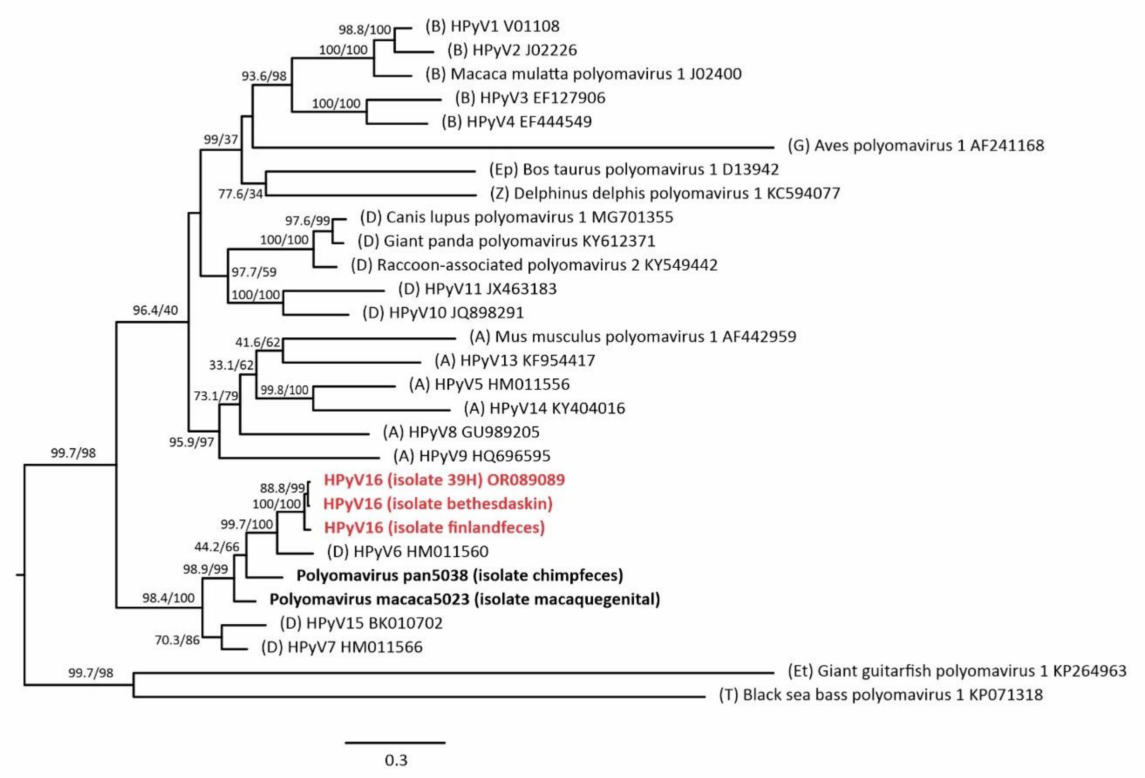


**Figure S1.** Maximum Likelihood phylogenetic tree (mid-point rooted) based on LT protein sequences for HPyV16 (in bold and red) and representative members of all genera of the *Polyomaviridae* family. The two distant HPyV6 relatives detected in macaque genital (isolate: macaquegenital) and chimpanzee fecal (isolate: chimpfeces) samples are also included (in bold). Alignment was built using Aliview [1] and conserved (non-ambiguous) sites were selected with GBlocks [2], in SeaView v.5 [3]. The analysis was performed under the LG+F+I+G4 model, estimated with ModelFinder in IQ-TREE [4]. The SH-aLRT/UFB values are indicated at nodes. The first letter/s in the sequence names indicates the genus (A): *Alphapolyomavirus*, (B): *Betapolyomavirus,* (D): *Deltapolyomavirus*; (Ep): *Epsilonpolyomavirus*, Et): *Etapolyomavirus,* (G): *Gammapolyomavirus*, (T): *Thetapolyomavirus*, (Z): *Zetapolyomavirus.*

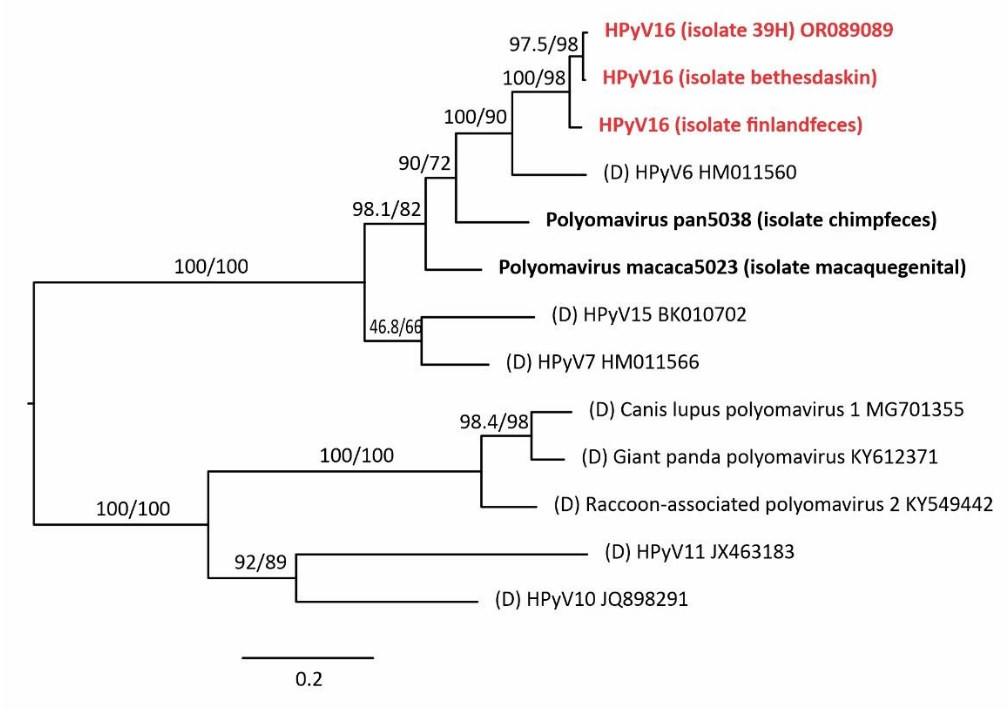


**Figure S2.** Maximum Likelihood phylogenetic tree (mid-point rooted) based on LT protein sequences for HPyV16 (in bold and red) and all members of the *Deltapolyomavirus genus (D).* The two distant HPyV6 relatives detected in macaque genital (isolate: macaquegenital) and chimpanzee fecal (isolate: chimpfeces) samples are also included (in bold). Alignment was built using Aliview [1] and conserved (non-ambiguous) sites were selected with GBlocks [2], in SeaView v.5 [3]. The analysis was performed under the LG+I+G4 model, estimated with ModelFinder in IQ-TREE [4]. The SH-aLRT/UFB values are indicated at nodes.

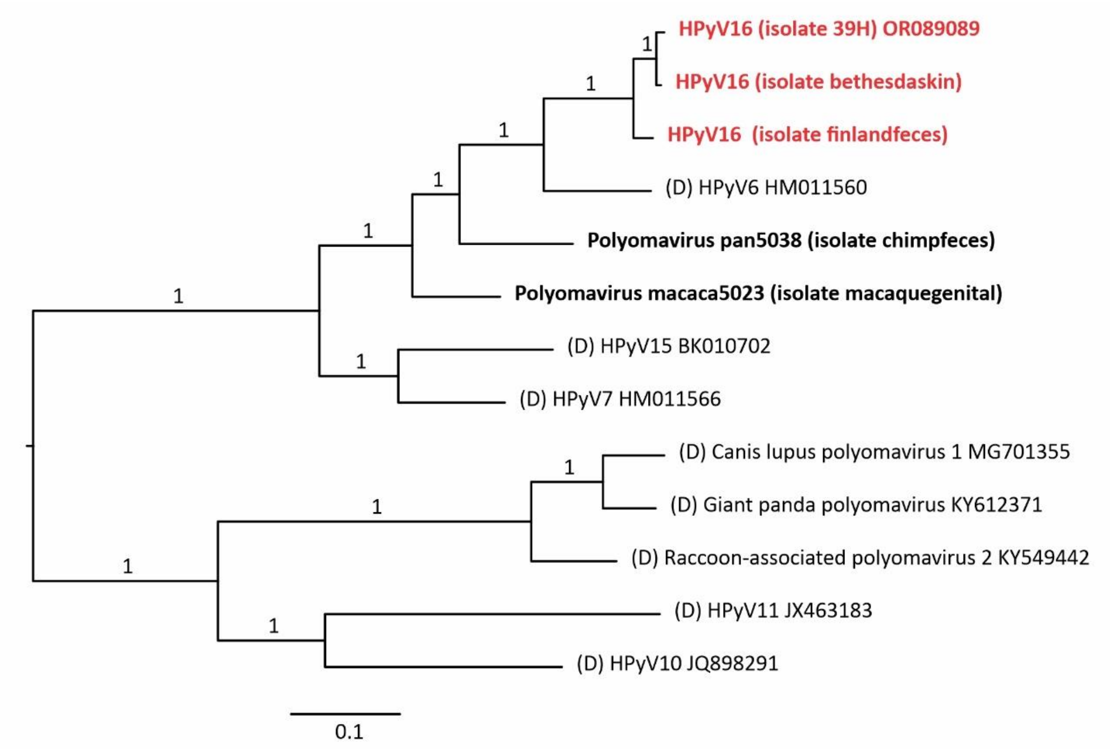


**Figure S3.** Majority-rule consensus tree (mid-point rooted) obtained in a Bayesian analysis based on LT protein sequences for HPyV16 (in bold and red) and all members of the *Deltapolyomavirus genus (D).* The two distant HPyV6 relatives detected in macaque genital (isolate: macaquegenital) and chimpanzee fecal (isolate: chimpfeces) samples are also included (in bold). Alignment was built using Aliview [1] and conserved (non-ambiguous) sites were selected with GBlocks [2], in SeaView v.5 [3]. The analysis was performed under the mixed model (where the Markov chain samples each model according to its probability) in MrBayes v3.2.7 [5]. The posterior probability is indicated at nodes.

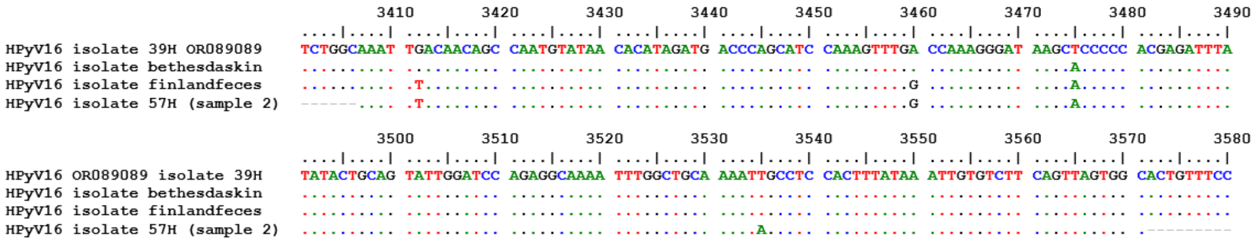


**Figure S4.** Alignment of HPyV16 sequences in the small region that could be obtained for the isolate 57H from a skin sample (sample 2) from Argentina (165 nts, positions 3407 to 3571 regarding the start of the LT of the HPyV16 isolate 39H (GenBank Accession Number: OR089089).
