## Supplementary Tables S1-S3 for "Identification of a new human polyomavirus in distinct populations and tissues"

### Supplementary material

**Table S1.** Primers and reaction condition for the amplification of complete genome sequences of HPyV16.

<table>
<tr><th>Reaction</th><th>Primers (5’ to 3’)</th><th>Conditions</th></tr>
<tr><td>RCA</td><td>ProS3: ATGTGGGAGGC<br>ProAS4: TGGKGCCATCTTCA<br>ProS5: AGRTAYTWTGG<br>ProAS6: HTTAAAATATCT<br>ProS7: GGVCARCCTATG<br>ProAS8: ATWYTDCAAAGTGG<br>ProS9: TTBACATCYTCAA<br>ProAS10: TTTAATTTGTAA<br>ProS11: WAYTAAWTTCCA<br>ProAS12: CWDTGGTTTGG<br>ProS13 CCTTTRTCWGGGTG<br><br>All primers have two 3'-terminal phosphorothioate modifications that make them resistant to the 3'→5' exonuclease activity.</td><td>RCA was performed using 5.0 μl of DNA extract, 10 U of phi29 DNA Polymerase (Thermo Scientific), 1X Rection Buffer, primers at a final concentration of 1 μM each, Non-acetylated BSA at a final concentration of 0.2 mg/ml, 2 mM dNTPs and nuclease free $H_2O$ in a final volume of 20 μl.<br><br>Reaction was incubated 18 h at 30 ºC, with an inactivation step of 10 min at 65 ºC.</td></tr>
<tr><td>PCR I</td><td>First round<br>S-BP2: CCTCAATGGMTGCTTTTT<br>H6-AS6: CTCATCTGGTGGTAAGTGC<br><br>Second round<br>S-BS2: ARCTTCCCAGAGTRATAA<br>H6-AS4: CAGGTRGTGATGCATTGAA</td><td rowspan="2">PCRs were performed using 4.0 ul of DNA from RCA (for the first round) or 2.0 ul of the first-round product (for the second round), 2 U of Expand™ Long Template PCR System (Roche), 1X ELT Buffer 2, primers at a final concentration of 0.4 uM each, 0.5 mM dNTPs and nuclease free H2O in a final volume of 25 ul.<br><br>Cycling conditions were: 2 min at 94 ºC, 10 cycles of 30 sec at 94 ºC, 30 sec at 45ºC (-0.5ºC/cycle), 2.5 min at 68 ºC, and 26 cycles of 30 sec at 94 ºC, 30 sec at 40ºC, 2.5 min at 68 ºC, and a final extension of 10 min at 68 ºC.</td></tr>
<tr><td>PCR II</td><td>First round<br>H6-S1: CCTTCTTTGTGCTGCTACTC<br>AS-BP2: TCTAWAGGCYTCCCAAAC<br><br>Second round<br>H6-S3: GGACTGCCATATTGTGGAG<br>AS-BS2: GCMTCAGGAATTTCAGGC</td></tr>
</table>

**Table S2.** Human polyomavirus numbering naming.

| Designation | Original name | Inferred Host | Associated disease | Citations |
|---|---|---|---|---|
| **HPyV1** | BK | human | Kidney bladder brain saliva | [1] |
| **HPyV2** | JC | human | Kidney bladder brain saliva | [2] |
| **HPyV3** | Karolinska Institute | human | - | [3] |
| **HPyV4** | Washington University | human | Pneumonia | [4] |
| **HPyV5** | Merkel cell | human | Skin carcinoma | [5] |
| **HPyV6** | HPyV6 | human | Pruritus | [6] |
| **HPyV7** | HPyV7 | human | Pruritus | [6] |
| **HPyV8** | Trichodysplasia spinulosa | human | TS | [7] |
| **HPyV9** | HPyV9 | human | - | [8] |
| **HPyV10** | Malawi, HPyV10 | human | - | [9,10] |
| **HPyV11** | Saint Louis | human | - | [11] |
| **HPyV12** | HPyV12 | Sorex araneus | - | [12] |
| **HPyV13** | New Jersey | human | Endothelial myositis | [13] |
| **HPyV14** | Lyon-IARC | Felis catus | - | [14] |
| **HPyV15** | Quebec | human | - | [15] |
| **HPyV16** | Finland | human | - | Current report |

[13] Mishra N, Pereira M, Rhodes RH, An P, Pipas JM, Jain K, et al. Identification of a Novel Polyomavirus in a Pancreatic

Transplant Recipient With Retinal Blindness and Vasculitic Myopathy. J Infect Dis 2014:1–5. https://doi.org/10.1093/infdis/jiu250.

[14] Gheit T, Dutta S, Oliver J, Robitaille A, Hampras S, Combes JD, et al. Isolation and characterization of a novel putative human polyomavirus. Virology 2017;506:45–54. https://doi.org/10.1016/j.virol.2017.03.007.

[15] Ondov BD, Starrett GJ, Sappington A, Kostic A, Koren S, Buck CB, et al. Mash Screen: high-throughput sequence containment estimation for genome discovery. Genome Biol 2019;20:232. https://doi.org/10.1186/s13059-019-1841-x.

**Table S3.** Splicing sites for HPyV6 and HPyV16.

| Polyomavirus | T antigen | Splice donor site (rating)[a,b] | Nt position | Splice acceptor site (rating) [a] | Nt position | Splice donor site (rating) [a] | Nt position | Splice acceptor site (rating) [a] | Nt position[c] |
|---|---|---|---|---|---|---|---|---|---|
| **Human polyomavirus 6 (HPyV6)** | 669T (LT) | GAGgtcagt (93) | 241-249 | tttattttacagGT (95) | 622-635 | | | | |
| | 190T (sT) | GTGgtaagt (93) | 567-575 | tttattttacagGT (95) | 622-635 | | | | |
| | 45T | GGGgtaatt (81) | 71-79 | aatgttttaaagGA (77) | 166-179 | GAGgtcagt (93) | 241-249 | tgatttcaatagAG (77) | 684-697 |
| **Human polyomavirus 16 (HPyV16)**[c] | 670T (LT) | GAGgttagt (93) | 238-246 | ttgtctttacagGT (95) | 616-629 | | | | |
| | 188T (sT) | GTGgtaagt (93) | 561-569 | ttgtctttacagGT (95) | 616-629 | | | | |
| | 48T[d] | GGGgtaatt (83) | 71-79 | aatattttaaagGA (80) | 166-179 | GAGgttagt (93) | 238-246 | agatttcaatagAG (73) | 678-691 |

[a] Splice sites for HPyV6 were experimentally determined [1]. Sequences are shown in the LT and sT open reading frames.
[b] Rating values were calculated with the Human Splice Finder tool [2] for the two HPyV16 detected in skin.
[c] Nucleotide positions are numbered regarding the start of the LT and sT coding sequences of the two HPyV16 detected in skin.
[d] Hypothetical protein here listed because of its close relationship to HPyV6.
